# An oral fentanyl self-administration model reveals dissociable escalation and relapse phenotypes in outbred vs inbred mice

**DOI:** 10.64898/2026.07.30.741697

**Authors:** Todd R. Appleby, Lukas K. MacMillen, Erica Sanchez, Sierra Schleufer, John F. Neumaier, Sam A. Golden, Kevin R. Coffey

## Abstract

Fentanyl-related overdose deaths now commonly involve non-injection routes, yet preclinical opioid self-administration is modeled predominantly intravenously. Here we establish an oral fentanyl self-administration procedure in male and female inbred C57BL/6 and outbred CD1 mice that measures volitional intake, cue-driven seeking, extinction, and relapse. Mice self-administered oral fentanyl (70 µg/mL) on a fixed-ratio 1 schedule across fifteen 3-hour sessions, followed by ten extinction sessions and a cued reinstatement test. A separate cohort underwent between-session dose thresholding across a quarter-log series from 222 to 22 µg/mL. Seventy-five percent of mice acquired self-administration, with similar rates across genetic background and sex. Responding increased as fentanyl concentration fell, indicating dose-sensitivity toward a preferred drug level. C57BL/6 mice escalated intake and lever pressing across sessions, responded persistently early in extinction before declining, and reinstated pressing to a conditioned cue. CD1 mice consumed high levels from the outset with limited escalation and showed neither extinction nor cued reinstatement of pressing, but shortened their reward-port approach latency when cues returned. This shows that lever presses alone would have misclassified them as weakly conditioned. A composite severity score summing seven components of fentanyl-use risk varied continuously rather than splitting into high and low groups, even among inbred mice. Sex differences were largely confined to C57BL/6 mice, in which females showed stronger cue association and higher severity scores than males. These results reveal separable escalation-prone and relapse-prone phenotypes that track genetic background. Protocols, hardware specifications, and analysis code are openly available, lowering the barrier to adopting oral fentanyl self-administration.

## Introduction

Opioid use disorder represents a significant public health crisis in the United States, with fentanyl in particular contributing to more than half of the ∼70,000 overdose-related fatalities in 2025^1^. Fentanyl is now the drug of choice for most opioid users, particularly for younger individuals and those suffering psychological distress^2^. Fentanyl induces rapid and short-lived euphoria followed by severe physical and emotional distress during withdrawal. In abstinence, fentanyl-related cues and hypersensitivity to emotional stressors motivate tenacious and repeated relapses^3–5^. Fentanyl also poses a unique risk compared to other opioids due to its extreme potency, low cost, and ease of administration. Currently approved treatments for fentanyl use disorder are similar to those used for other opioids, primarily opioid replacement therapies or opioid receptor blockade^6^. These strategies have unfortunately proven to be insufficient to stem the tide of fentanyl-related suffering and death. Accessible and reproducible preclinical models that capture volitional fentanyl use remain essential to developing the next generation of treatments.

Preclinical opioid self-administration is modeled predominantly by the intravenous route, yet non-injection routes now predominate among human overdose deaths^7,8^. Oral ingestion is especially relevant given the prevalence of illicit fentanyl tablets in the United States^9^. Existing mouse models deliver fentanyl intravenously^10,11^, by vapor^12,13^, or in home-cage self-administration contexts^14^. Similarly, oral consumption has been examined under aversion-resistance oral home-cage and self-administration procedures^15^. No oral procedure in mice currently resolves lever approach, cue presentation, reward-port approach, and consumption as separate events across acquisition, extinction, and cued reinstatement, as we and others have described in rats using oral fentanyl^16^ and oxycodone^17^.

Opioid use disorder (OUD) is complex and difficult to model preclinically. A major facet of OUD that is often understudied is the substantial individual differences in the severity of opioid use and relapse, along with the behavioral predictors that distinguish vulnerable from resilient individuals. Individual differences of this kind are well established in preclinical reward research, where animals differ reliably in the incentive-motivational value they assign to reward cues^18,19^ and multi-criteria approaches have been used to rank animals along several dimensions of drug-seeking severity at once^16,20,21^. To address this, we developed a robust and convenient fentanyl self-administration (SA) model that 1) measures key components of opioid use such as intake, seeking, extinction, and reinstatement, and 2) uses these measures for studying individual differences in risk and resilience toward volitional fentanyl intake and reinstatement in male and female mice. To facilitate adoption across laboratories, we provide an open community resource containing detailed protocols for implementing the model with relatively low-cost, widely available equipment, together with code for standardized analysis of operant measures ^22^

On a group level, mice in this procedure show the classical features of volitional drug intake: three quarters acquired fentanyl self-administration, responding came under the control of conditioned cues, and seeking returned when those cues were presented after extinction. Both strains increased responding as fentanyl concentration fell, indicating adjustment toward a preferred drug level rather than a fixed response habit. Because lever pressing, cue presentation, reward-port approach, and consumption are resolved as separate events, the procedure exposes differences that any single measure would obscure. Inbred C57BL/6 mice escalated intake and reinstated lever pressing, whereas outbred CD1 mice consumed high levels from the outset without escalating and expressed cue-evoked seeking as faster reward-port approach rather than lever pressing. Sex differences were limited and largely confined to C57BL/6 mice, and severity varied continuously within both strains, indicating that genetic background alone does not account for individual vulnerability. These unique features provide valuable guidance for researchers interested in studying specific components of opioid use disorder in mice.

Overall, this model provides an adaptable behavioral entry point for studying fentanyl seeking, onto which the genetic, physiological, and brain-wide imaging tools available in mice can be brought to bear.

## Results

### Oral Fentanyl Self-Administration is Acquired by Male & Female C57 and CD1 Mice

All mice that completed the standard SA protocol were used to determine acquisition rates for C57BL/6 and CD1 mice (Figure 1a). Acquisition was defined as earning an average of 10 rewards per session during SA training sessions 6-15 (Figure 1b). Seventy-five percent of all mice acquired SA, with C57BL/6 mice acquiring at a slightly higher rate (80%) compared to CD1 mice (70%, Figure 1c). Acquisition rates were nearly identical for males and females within each strain. *Earned Rewards* increased over sessions as evidenced by a significant main effect of *Session* (β= 0.8, SE= 0.2, t(1225)= 3.8, p< 0.001). Though C57BL/6 and CD1 mice eventually earned equivalent rewards per session, there was both a significant main effect of *Strain* (β= -15.6, SE= 4.6, t(1225)= -3.4, p< 0.001), and a significant *Strain x Session* interaction (β=1.5, SE=0.3, t(1225)= 5.5, p< 0.001). These effects reflect the different ways each strain learns to SA, with CD1 mice earning many rewards early and increased intake gradually, while C57BL/6 mice start with very few rewards and escalate at a steeper slope (Figure 1d).

**Figure 1.**
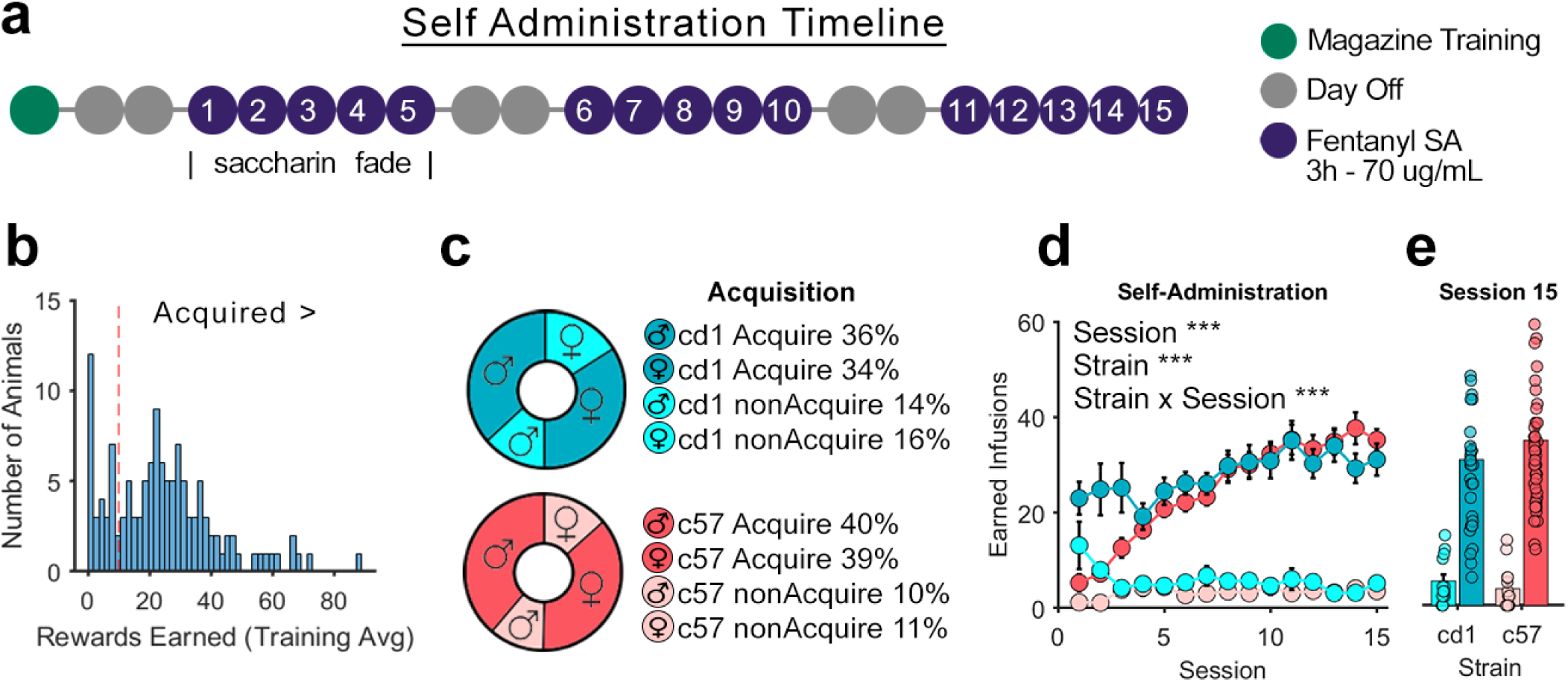
Acquisition of Oral Fentanyl Self-Administration. **a)** Timeline for all self-administration experiments. **b)** The overall distribution of average rewards earned during training was used to determine the acquisition threshold (Avg > 10). **c)** Acquisition rates for c57 and CD mice were similar (80-70% respectively), and males and females were very similar withing strain. **d)** c57 and CD1 mice show different self-administration trajectories, with c57 mice learning to escalate over session, while CD1 mice immediately begin and maintain high intake. **e)** by Session 15, infusions earned are similar across strains. **P<0.01

### C57BLc & CD1 Mice Display Unique Behavioral Trajectories during Self-Administration, Extinction, and Reinstatement Testing

Male and female C57BL/6 and CD1 mice underwent 3 weeks of oral fentanyl self-administration, followed by 2 weeks of extinction and a single (1 h) cued reinstatement testing session (Figure 2a). Male and female C57BL/6 mice began with low intake (∼0.5 mg/kg) and significantly escalated fentanyl intake across sessions (β= 161.8, SE= 29.7, t(494)= 5.4, p< 0.001; Figure 2b). In contrast, male and female CD1 mice began training with relatively high intake (∼1.5 mg/kg) but showed no significant change in intake across sessions (Figure 2c). This pattern was mirrored in active lever presses, where C57BL/6 mice showed a significant increase over training (β= 3.7, SE= 0.7, t(494)= 5.5, p< 0.001; Figure 2d), while CD1 mice did not (Figure 2f). C57BL/6 mice then showed a significant decrease in active lever pressing during extinction (β= -4.4, SE= 2.1, t(324)= - 2.1, p= 0.037; Figure 2e), followed by significant cued reinstatement of lever pressing (β= 3.7, SE= 1.5, t(64)= 2.5, p= 0.015; Figure 2e). In contrast, CD1 mice did not show significant extinction of lever pressing or evidence of cued reinstatement of lever pressing (Figure 2g). C57BL/6 mice also showed a small but significant increase in inactive lever pressing during SA training (β= 1, SE=0.4, t(494)= 2.7, p= 0.007; Figure 2h), though the slope of this increase was much lower than for active presses (1 vs 3.7). No other differences in inactive lever pressing were observed (Figures 2i,j,k). Head entry latency into the reward delivery cup after an active lever press was measured to assess cue-reward association. There was no main effect of *Session* on head entry latency for C57BL/6 mice, likely because head entry latency dropped almost immediately on session 2 and remained low throughout SA training for females and slightly increased again for males. This pattern caused a linear fit with minimal slope (β= -1). By session 15, female C57BL/6 mice had significantly lower head entry latency than males (β= 26.4, SE= 8.2, t(32)= 3.2, p= 0.003; Figure 2l). Head entry latency then significantly increased for C57BL/6 mice during extinction, when cues were not present (β= 2.8, SE= 0.6, t(324)=4.6, p< 0.001; Figure 2m), with male C57BL/6 mice showing higher latency than females (β= 41.6, SE= 16.9, t(324)= 2.5, p= 0.014; Figure 2m). C57BL/6 mice did not show a significant reduction in head entry latency during cued reinstatement testing. On the other hand, CD1 mice showed a significant decrease in head entry latency during SA training (β= -2.1, SE= 0.7, t(210)= -2.9, p= 0.005; Figure 2n), followed by a significant increase in latency during extinction (β= 3.2, SE= 1, t(139)=3.2, p= 0.002; Figure 2o). CD1 mice then showed a significant decrease in head entry latency during cued reinstatement testing (β= -38.2, SE= 12.7, t(26)= -3, p= 0.006; Figure 2o). Together these data indicate substantial sex and strain differences in self-administration, extinction, and reinstatement. In particular, C57BL/6 mice show significant escalation of intake while CD1 mice show substantial but consistent intake. Further, extinction and reinstatement may be better captured by lever pressing in C57BL/6s and by head entry latency in CD1 mice and may reflect different features in learned reward seeking behaviors. These differences are further explored in our fentanyl risk and resilience analysis.

**Figure 2.**
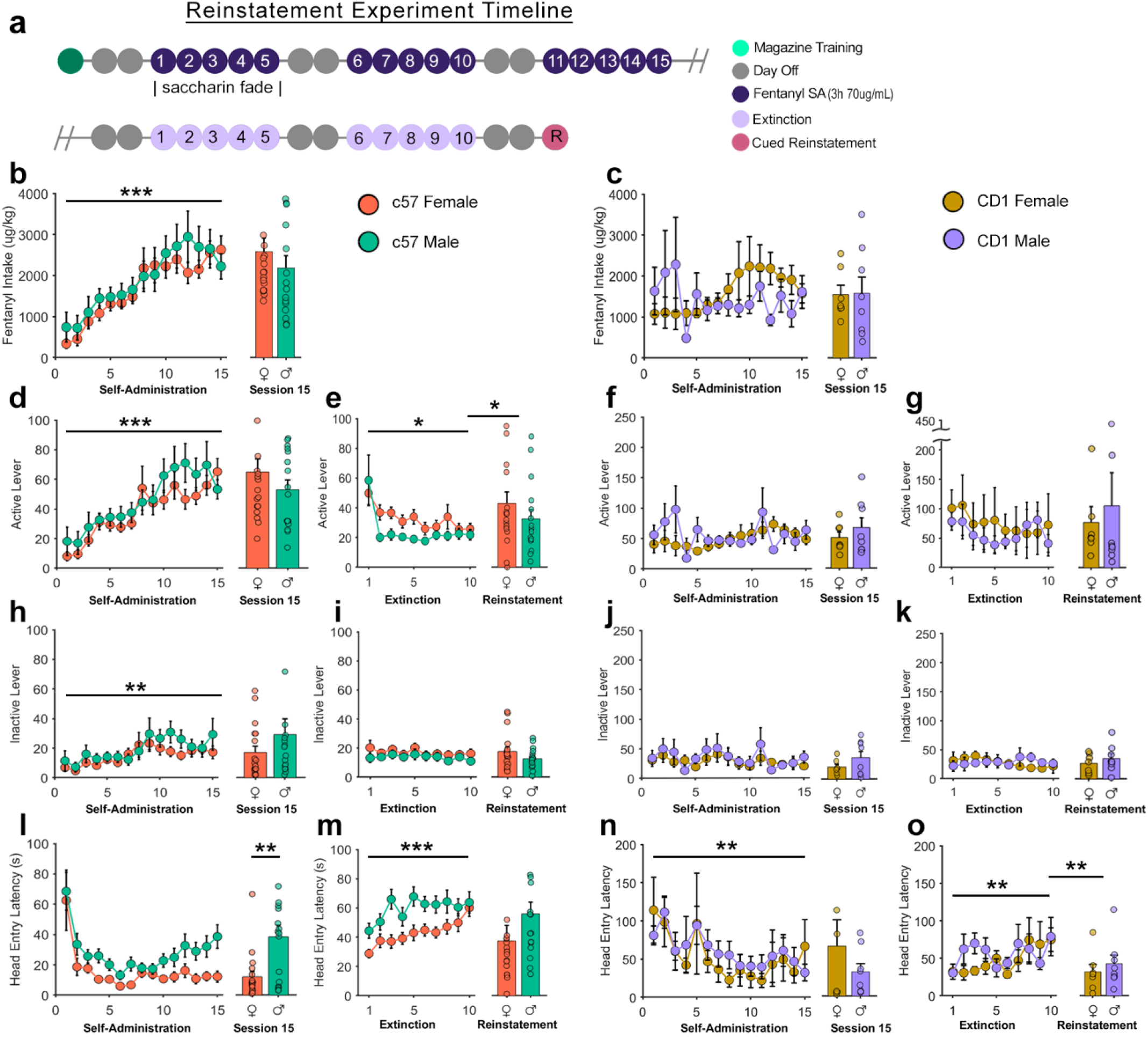
Oral Fentanyl Self-Administration, Extinction, & Reinstatement. **a)** Timeline for cued-reinstatement experiment. Daily fentanyl intake across SA sessions for **b)** c57 and **c)** CD1 mice. Daily active lever pressing trajectories for **d)** c57 mice during SA, **e)** c57 mice during extinction and reinstatement, **f)** CD1 mice during SA, and **g)** CD1 mice during extinction and reinstatement. Daily inactive lever pressing trajectories for **h)** c57 mice during SA, **i)** c57 mice during extinction and reinstatement, **j)** CD1 mice during SA, and **k)** CD1 mice during extinction and reinstatement. Average head entry latency after lever pressing for **l)** c57 mice during SA, **m)** c57 mice during extinction and reinstatement, **n)** CD1 mice during SA, and **o)** CD1 mice during extinction and reinstatement. ***P<0.001, **P<0.01, *P<0.05

### Mice Increase Responding to Decreasing Doses of Fentanyl in Order to Maintain Relatively Stable Intake

A separate cohort (n=12 in each group) of male and female CD-1 mice and male and female C57BL/6 mice underwent a between-sessions thresholding procedure to explore dose-intake characteristics. Similar to the extinction and reinstatement experiment, C57BL/6 mice displayed a significant increase in fentanyl consumption (β= 170.6, SE= 23, t(281)= 7.4 p< 0.001; Figure 3b) and active lever pressing (β= 4.4, SE= 0.8, t(281)= 5.3 p< 0.001; Figure 3f) over sessions, while CD1 mice maintained stable intake and pressing across sessions (Figure 3d,h). After 3 weeks of SA, animals underwent a 5 day thresholding procedure where dose began high and decreased each day according to a (¼)Log series (222, 125, 70, 40, 22 μg/mL). Intake and responding on each session were compared to the 70 µg/mL session, as it was expected concentration of fentanyl.

**Figure 3.**
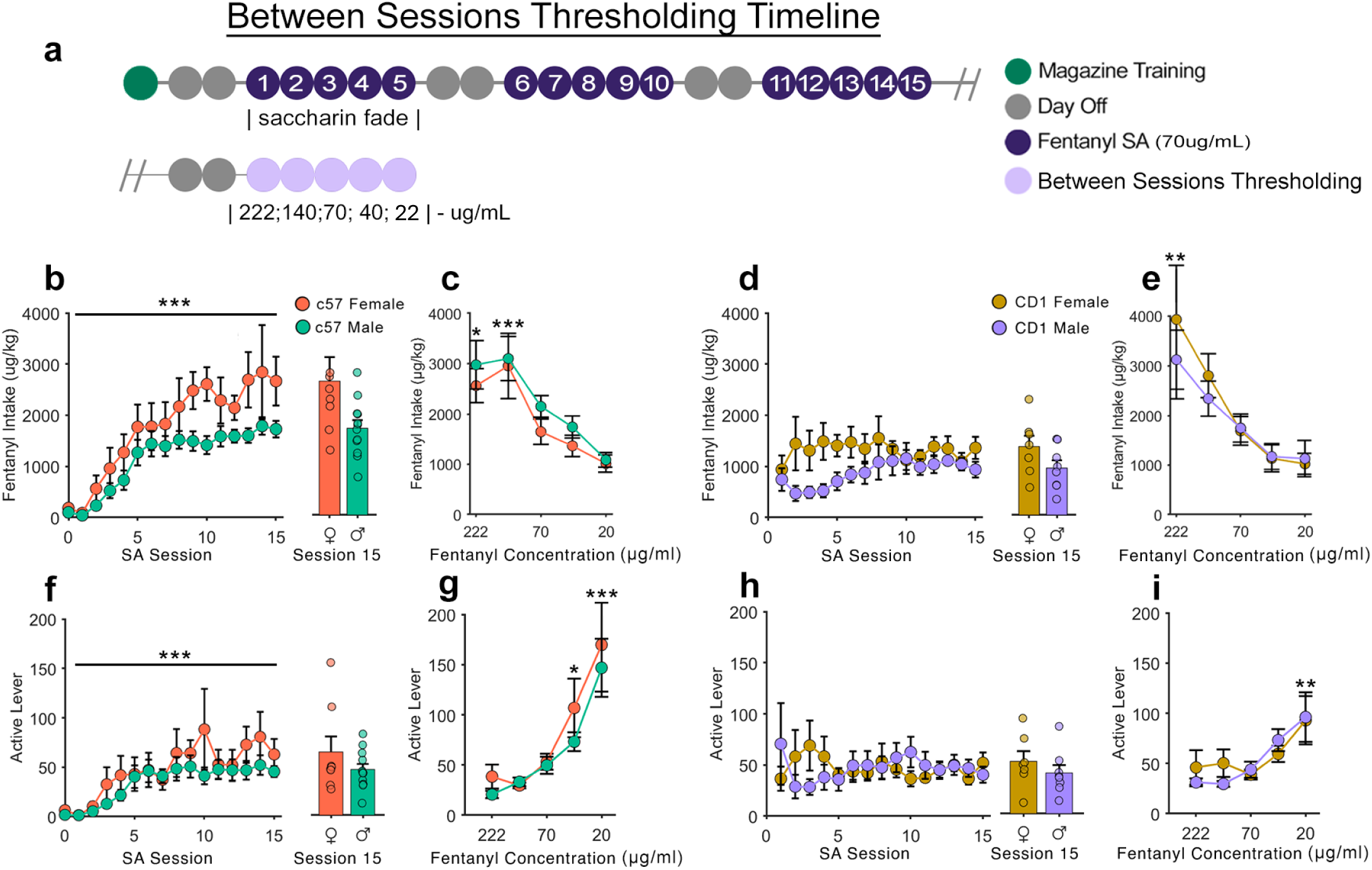
Between Sessions Thresholding. **a)** Timeline for the between-sessions thresholding experiment. **b)** Daily fentanyl intake across SA sessions (estimated from earned rewards and head entries) for c57 mice**. c)** Measured fentanyl intake across sessions with differing fentanyl concentrations for c57 mice. **d)** Daily fentanyl intake across SA sessions (estimated from earned rewards and head entries) for CD1mice**. e)** Measured fentanyl intake across sessions with differing fentanyl concentrations for CD17 mice. **f)** Active lever presses across sessions for c57 mice. **g)** Active lever presses across sessions with differing fentanyl concentrations for c57 mice**. h)** Active lever presses across sessions for CD1 mice. **i)** Active lever presses across sessions with differing fentanyl concentrations for CD1 mice. For SA sessions, ***P<0.001 (linear effect of session). For Between Sessions Thresholding, ***P<0.001, **P<0.01, *P<0.05 (categorical effect of session as compared to 70ug/mL session).

During IV self-administration, animals can maintain stable drug intake across a wide range of doses due to their ability to rapidly detect changes in drug level^23^. However, this is more difficult in an oral SA procedure where the time between consumption and sensation is extended. Accordingly, C57BL/6 mice consumed significantly more fentanyl on the high dose sessions of 222 µg/mL (β= 915.8, SE= 362.5, t(85)= 2.5, p= 0.013; Figure 3c), and 125 µg/mL (β= 1308.3, SE= 362.5, t(85)= 3.6, p< 0.001; Figure 3c), while CD1 mice consumed significantly more fentanyl on the 222 µg/mL session (β= 2259, SE= 743.5, t(66)= 3, p= 0.003; Figure 3i). However, these mice were still clearly able to detect the falling concentrations of fentanyl, with C57BL/6 mice significantly increasing active lever pressing on the low concentration 40 µg/mL (β= 53.6, SE= 25.7, t(85)= 2.1, p= 0.04; Figure 3g) and 22 µg/mL sessions (β= 116.5, SE= 25.7, t(85)= 4.2, p< 0.001; Figure 3g), while CD1 mice significantly increased active lever pressing on the 22 µg/mL session (β= 46.1, SE= 17.2, t(85)= 2.7, p= 0.009; Figure 3g). Overall, C57BL/6 mice were more sensitive to changing drug concentrations, and more flexible in changing their response rates.

### Oral Fentanyl Self Administration in Mice has Distinct Loading and Maintenance Phases

Similar to our previous work in Long-Evans rats^16^, oral fentanyl SA in mice appears to be composed of two distinct phases, a ‘loading’ phase where animals rapidly consume drug to reach a preferred drug level, followed by a ‘maintenance’ phase where that level is maintained with slower, evenly spaced consumption. This pattern can be seen in the estimated brain fentanyl concentration of C57BL/6 (Figure S1a,b,c) CD1 (Figure S1d,e,f) along with their corresponding cumulative response distributions Figure S1g,h,i,j,k,l). While the preferred drug level of CD1 mice remained similar across weeks, C57BL/6 tended to escalate their preferred level. C57 mice accomplished this by increasing the number and rate of operant responses during both the loading phase and the maintenance phases in later sessions as training proceeds. Because of the exponential nature of pharmacokinetics, maintaining a higher drug level requires an increase in response rate, as evidenced by steeper cumulative response function during the maintenance phase on later sessions (Figure S1g vs Figure S1h,i).

### C57BL/c and CD1 Mice Display Individual Differences in Fentanyl Use Risk and Resilience

A descriptive analysis of individual differences in multiple behaviors was performed to ascertain the underlying behavioral distribution of fentanyl use risk severity. Although they are inbred, C57BL/6 mice still display wide distributions of behavior across multiple domains of fentanyl use risk, including Intake, Seeking, Escalation, Extinction, Recall, and Relapse (Figure 4a-h). By summing each animal’s normalized risk for each behavioral component, we can provide a composite severity score that showed that most female C57BL/6 mice expressed greater risk characteristics than males, while they also had greater outliers with very high and very low risk metrics (Figure 4h). Outbred CD1 mice also showed a wide range of behaviors across these components (Figure 4i-o), with some notably high outliers in extinction and relapse responding. Composite severity scores for CD1 mice were relatively evenly spread around 0 (Figure 4p). Cross correlations were performed to determine if particular risk components were correlated within strain or sex. For C57BL/6 males and females, along with CD1 females, total intake was highly correlated with escalation (Figure 4q,r,s), but for CD1 males, they were not (Figure 4t). This reflects the propensity for some CD1 males to begin self-administering high quantities immediately, but to then fail to escalate. For CD1 mice, extinction and relapse responding were highly correlated, reflecting their strong persistence in lever pressing regardless of drug and cue availability (Figure 4t). PCA analysis was also used to visualize the contribution of different risk components to each animal’s overall severity profile (Figure 4u-w). These figures highlight the different behavioral profiles for each strain. High risk C57BL/6 animals seem particularly driven by high intake, escalation and seeking (Figure 4u,v), while high risk CD1 animals are better defined by persistence in extinction responding, relapse, and recall (Figure 4w). This could be highly valuable information for determining the optimal animal model for researchers interested in studying escalation or high drug intake (C57BL/6s) vs those interested in studying the mechanisms of relapse (CD1) and are likely to reflect distinct patterns of regional neural plasticity that develop during opioid SA between strains.

**Figure 4.**
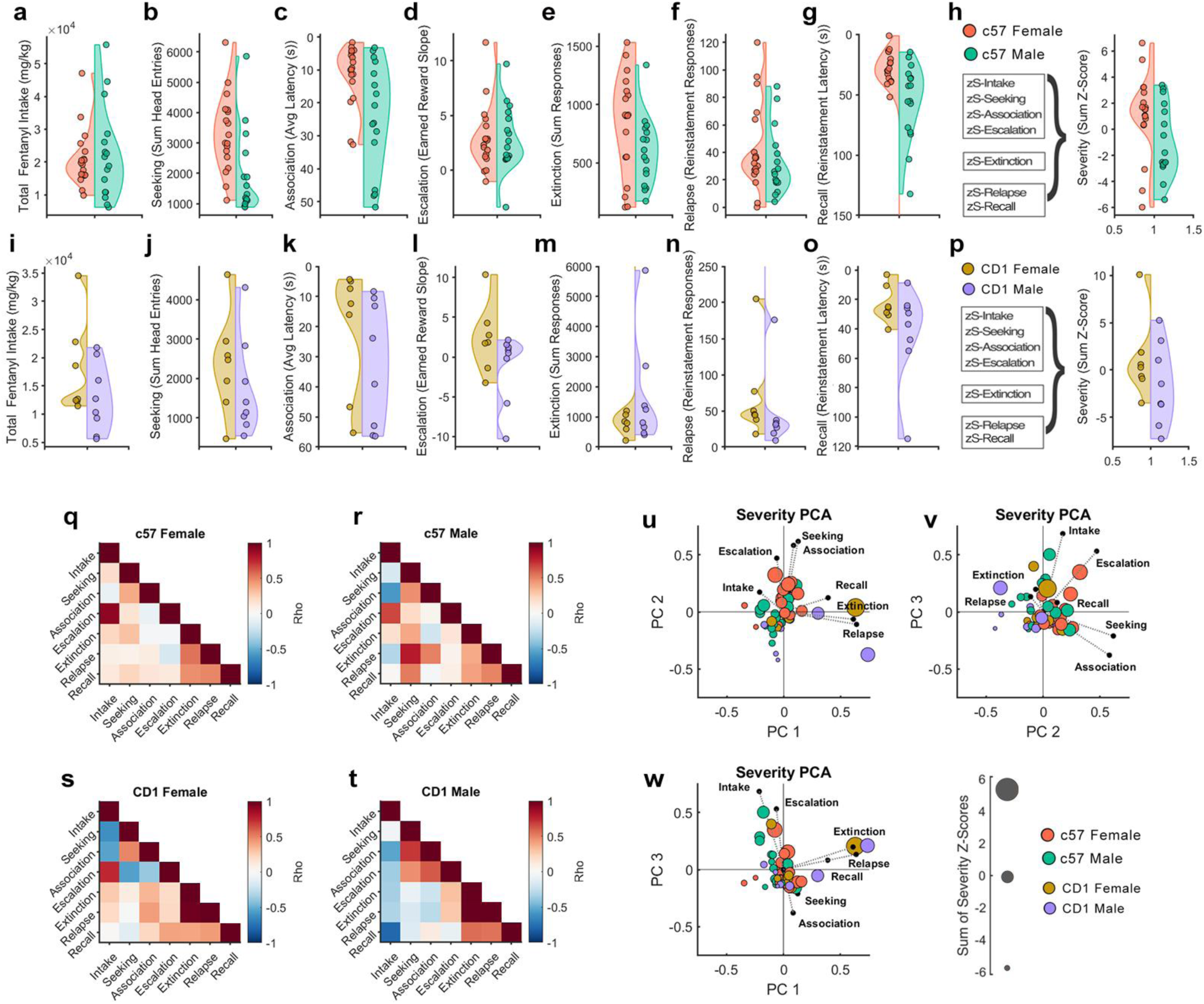
Individual Differences in Fentanyl Risk Severity. Behavioral distributions for c57 mice for unique components of OUD such as **a)** total fentanyl intake, **b)** fentanyl seeking, **c)** cue association, **d)** escalation, **e)** extinction responding, **f)** relapse responding, **g)** cue recall and **h)** composite severity score. Behavioral distributions for CD1 mice for unique components of OUD such as **i)** total fentanyl intake, **j)** fentanyl seeking, **k)** cue association, **l)** escalation, **m)** extinction responding, **n)** relapse responding, **o)** cue recall and **p)** composite severity score. Operational definitions for each component are provided in the methods section. Fentanyl risk component cross-correlations for **q)** c57 females, **r)** c57 males, **s)** CD1 females, and **t)** CD1 males. **u, v,w)** Principal component analyses to visualize the contribution of each risk component to individual animal composite risk score.

## Methods

### Animals

For all experiments, 7-12 week-old male and female CD-1 (Charles River) and C57BL/6 (bred in-house) mice were used. Mice were group-housed in a temperature and humidity-controlled room on a 12-hour reverse dark-light cycle and acclimated to the facility and handling for at least 1 week before beginning experimental procedures. Food and water were available ad-lib throughout the experiment (weight range: 15-45g). Experimental procedures were approved by the University of Washington Institutional Animal Care and Use Committee.

### Experimental Design: Oral Fentanyl Self-Administration, Extinction, & Reinstatement Testing

Male (n=10) and female (n=10) CD-1 mice (Charles River) and male (n=21) and female (n=22) C57BL/6 mice were trained to self-administer (SA) liquid fentanyl. Mice were first exposed to the operant chamber during a 1 hr magazine training session (Friday) where 1 reward/cue is unconditionally administered every 60 s. The audiovisual conditioned stimulus (CS) consisted of illumination of a yellow light above the active lever for 1-second coincided with a 5-second tone (4 kHz) and was paired with delivery of a 0.1% saccharin reward in HydroPak H2O (.02 mL). The following Monday, mice learned to SA fentanyl (70 μg/mL) in H2O (0.02 ml/delivery) for 5 days a week (3 hr/session) over three weeks on a fixed-ratio 1 schedule (Figure 1a). The first 5 days of SA employed a “saccharin fade” (0.1%, 0.08%, 0.06% 0.04%, 0.02%) where the full dose of fentanyl was present from day 1. Upon active lever press, liquid fentanyl was delivered to a small dish paired with the above-described CS. During the CS presentation, a time-out was imposed. SA occurred during dark cycle in standard operant chambers (Med-Associates) with 2 levers and a low-cost IR USB camera (IMX462, Arducam). Presses on the inactive lever were recorded with no programmed consequences. On experimental day 22, mice began extinction training wherein lever pressing resulted in neither cue presentations nor fentanyl delivery. Extinction sessions (3 hr/session) continued for 10 days for all animals. Seventy-two hours after the final extinction session, animals underwent cued reinstatement testing (1 hr/session) where the compound CS signaled the beginning of the session. Each subsequent active lever press elicited the compound CS but did not result in the delivery of fentanyl. These procedures were modified from our previously published rat SA protocol^16^. A complete behavioral protocol and description of the chambers can be found in our [code and data repository](https://github.com/schleuf/CoffeyBehavior_MouseEdition).

### Experimental Design: Between Session Thresholding

A separate cohort of male (n=12) and female (n=12) CD-1 mice and male (n=12) and female (n=12) C57BL/6 mice were used to further validate our oral-fentanyl SA model with a between-sessions-thresholding procedure to explore dose-intake characteristics. Mice began SA training as described above. On the 11^th^ day of SA training, animals began the between-sessions-thresholding procedure. For this procedure, the concentration of fentanyl was progressively decreased each day for 5 days following a (¼)Log series (222 μg/mL, 125 μg/mL, 70 μg/mL, 40 μg/mL, 22 μg/mL). Fentanyl is absorbed rapidly through the oral mucosa allowing animals to assess changes in dose^24^. In support of this, animals did not always consume all the liquid that was delivered on high concentration days. However, because the liquid was delivered via a syringe pump to a deep dish where no spillage occurred, we were able to draw back the remaining liquid at the end of each session to measure total consumption precisely.

### Modeling Estimated Brain-Fentanyl Concentration

Whole-brain levels of fentanyl were estimated using a two-compartment model^25^. Briefly, we used the equation:

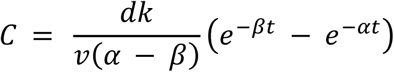

Here, C is the concentration in the brain (µg/kg), *d* is the dose, *k* is the rate constant for transfer from the body to the brain (0.233), *v* is volume of the brain (0.15 l/kg), *α* and *β* (0.642 and 0.0971, respectively) are constants representing the flow of fentanyl between the body and brain compartments and the elimination of fentanyl from the body, and *t* is the time in min since the last infusion. All constants were estimated using methodology derived from intraperitoneal (IP) cocaine experiments, but with a larger elimination coefficient (*kel=0.294)*. Using coefficients from IP as opposed to PO experiments was chosen because ∼50% of oral fentanyl is absorbed directly through the oral mucosa, while only half of the remaining 50% escapes first-pass metabolism in the gut^24^. Still, these values represent educated guesses and are subject to change upon measurement of oral fentanyl pharmacokinetics in mice.

### Statistical Analyses

Statistical analyses for oral fentanyl SA needed to account for repeated measures over time, between groups difference, some missing data, and nesting within individual animals. To accomplish this, all statistical analyses were run using linear mixed-effects modeling (LME). LMEs are flexible, robust, and able to be tailored to each experiment’s design and specific questions of interest. Complete statistical analyses and outputs can be viewed in our data repository, and statistics are reported in the manuscript for fixed effects coefficients in the form of:

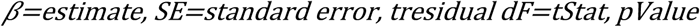

### Overall Self-Administration Acquisition

For the acquisition analysis the LME used fixed-effects terms for *Sex, Session,* and *Strain,* their interaction terms, and a random intercept that varied by *Subject.* LMEs were calculated with MATLAB’s fitlme() function with the following general formula:

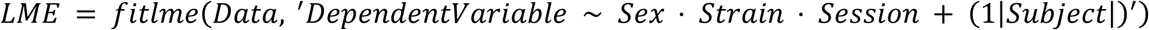

### Self-Administration Extinction and Reinstatement Experiment

For the extinction and reinstatement experiment, LMEs were run for each strain individually to explore sex and session differences within strain, and for sex differences on key days (session 15, and reinstatement). The Reinstatement session was also compared to hour 1 of extinction session 10, to determine if the reintroduction of cues reinstated different behaviors. The comparator group was C57BL/6 for *Strain*, Male for *Sex,* while *Session* was modeled linearly. Here, we also included *Session* as a random variable allowing the model to use a unique intercept and slope for each subject to account for individual differences in starting behavior and learning rate. LMEs were calculated with MATLAB’s fitlme() function with the following general formulas:

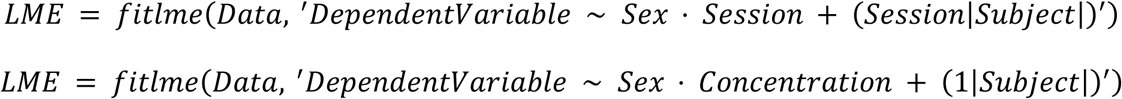

### Oral Fentanyl SA Individual Differences & Risk Analysis

A descriptive analysis of individual differences in multiple behaviors was performed to ascertain the underlying behavioral distribution of fentanyl use risk severity. Different components of substance use disorder were operationalized using behaviors from our model to improve interpretability. *Intake* was defined as the sum of fentanyl intake on SA sessions 6-15; *Seeking* was defined as the sum of head entries on SA sessions 6-15; *Cue Association* was defined as a maximum latency of 360s minus the average head entry latency after an active lever press on SA sessions 6-15 (low latency values represent high levels of association); *Escalation* was defined as the slope of earned rewards from sessions 1-10; *Extinction* was defined as the sum of active lever pressing during extinction sessions 1-10; *Relapse* was defined as the number of active lever presses during reinstatement testing; *Recall* was defined as a maximum latency of 360s minus the average head entry latency after an active lever press during the reinstatement session; finally the composite *Severity* score was defined as the sum of Z-Scores from all individual components. Cross correlations were performed to determine if particular risk components were correlated within strain or sex, and all SA animals were entered into a PCA analysis using all risk components as the input vector to visualize the contribution of different risk components to each animal’s overall severity profile.

### Code and Data Availability

All code and processed data used to generate figures and statistics for this manuscript are available on a GitHub repository: https://github.com/DrCoffey/CoffeyBehavior_MouseEdition and archived at Zenodo (doi:10.5281/zenodo.21694585).

## Discussion

The oral fentanyl self-administration procedure described here resolves intake, escalation, extinction, and cue-induced reinstatement in male and female C57BL/6 and CD1 mice, and our central result is that these components do not vary together. Mice of both strains and sexes acquired self-administration, formed reliable associations between the operant response and reward delivery, adjusted responding when fentanyl concentration changed, sustained responding early in extinction, and renewed pursuit of fentanyl when conditioned cues returned (Figs 1-3). Against that shared background, strain-dependent differences emerged in intake and active-lever trajectories. C57BL/6 mice began at low intake and progressively escalated consumption and active-lever responding, whereas CD1 mice established high intake early and maintained it across training. The strains also expressed conditioned fentanyl seeking differently. C57BL/6 mice showed conventional cue-induced reinstatement of lever pressing, whereas CD1 mice showed cue-sensitive changes in the latency to approach the reward receptacle. We find that Intake, Seeking, Cue Association, Escalation, Extinction, Relapse, and Recall are distinct indices of fentanyl-use risk severity that combined into separable fentanyl-seeking phenotypes differing largely based on genetic background rather than sex.

Our data indicates that escalation and high intake are dissociable aspects of fentanyl-seeking. Many CD1 mice consumed large quantities from the outset without subsequently escalating, whereas C57BL/6 mice began low and escalated steeply. Voluntary oral opioid intake in mice is under substantial genetic control, varying several-fold across inbred strains^26^ and mapping in part to a locus on chromosome 10^27^, so genetic background is one possible source of difference between strains of mice The strain difference could also arise from processes we did not measure, including tolerance, withdrawal severity, affective state, or motivational vigor. Because the procedure requires no surgery and supports prolonged longitudinal testing, each of these can be assayed in the same animals alongside the components reported here, and we regard them as predictions of this framework rather than equally likely alternatives to it.

In both CD1 and C57BL/6 mice, we show that mice will titrate their lever presses during self-administration based on fentanyl concentration (Fig. 3). For example, both strains increased active-lever responding at lower concentrations. This indicates that mice are seeking to reach a specific drug concentration in the brain, a phenomenon reflected in within-session model-based estimates of onboard fentanyl (Supplemental Fig. 1). These findings demonstrate concentration-sensitive adjustment and preferred drug state rather than a formal behavioral-economic measure of demand.

Oral self-administration characterizes fentanyl seeking by lever press, cue delivery, port entry, and consumption measures. How each of these measures were driven by the conditioned cue differed by strain. C57BL/6 mice reduced lever pressing during extinction and increased it when fentanyl-associated cues were restored. CD1 mice showed no corresponding change in pressing. Rather, their port-entry latency lengthened when cues were absent and shortened when cues were returned (Fig 2). Therefore, a measure of fentanyl seeking characterized by pressing alone would have classified CD1 mice as showing weak conditioned fentanyl seeking, when the same animals showed robust cue-evoked reward approach. CD1 lever pressing was in fact largely non-specific: extinction responding remained high and correlated with reinstatement responding, consistent with generalized motivation rather than cued seeking. Their reward-port approach behaved differently, lengthening as cues were withheld across extinction and shortening when cues were restored (Fig 2o). This within-subject reversal tracks cue availability and is not predicted by non-specific arousal, indicating that CD1 mice retained the cue-reward association and expressed it through reward approach rather than lever pressing.

Both strains showed broad individual variation across all severity score components. There were few consistent sex differences. C57BL/6 males were slower than females to approach the reward port during self-administration (Fig 2l), and there was not a consistent sex difference in fentanyl intake between experimental cohorts (Fig 2b and 3b). We therefore treat sex as unresolved in this model. Intake and escalation were strongly related in all groups except CD1 males. In C57BL/6 mice, females scored higher on average and spanned a wider range. High-scoring C57BL/6 mice were characterized primarily by intake, escalation, and seeking, whereas elevated CD1 profiles were associated with persistent pressing during extinction, reinstatement responding, and cue recall (Fig. 4). These analyses are descriptive, and several components share source data: Intake and Escalation are both derived from the reward-count series, and Cue Association and Recall are the same latency measure computed during self-administration and at reinstatement. We therefore interpret the component structure as separable behavioral profiles rather than statistically independent dimensions^11,14^. The relative lack of sex-specific effects suggests genetic background had a stronger pull on risk severity in our model. Quantitative dissociations of fentanyl risk taking factors have also been reported in other, non-oral self-administration models. For example, extended-access fentanyl vapor self-administration yields separable measures for intake/motivation and hyperalgesia/punished-seeking^13^.

Severity scores varied continuously within each strain and sex group, with no clear division between high- and low-risk animals. Notably, substantial variation was also present among inbred C57BL/6 mice, showing that genetic background alone does not account for all individual differences.

The procedure described here complements intravenous and vapor approaches^10,12,13^. It avoids surgery, supports prolonged longitudinal testing, produces robust responding without food or water restriction, and separates lever approaches, cue presentation, reward port approaches, and consumption as distinct measurable events. Together with the analysis tools provided here, our oral self-administration model can be easily adopted using widely available equipment.

In addition to the limitations described above, orally consumed fentanyl is absorbed more slowly and variably than intravenous or inhaled fentanyl, and direct pharmacokinetic measurements were not obtained. The brain-concentration curves in Supplemental Figure 1 depend on a two-compartment model with parameters taken from Pan et al. (1991)^25^ and on absorption assumptions drawn from Naji et al. (2023)^24^, neither established for oral fentanyl in mice, and should be read as model-based expectations rather than measurements. Withdrawal, progressive-ratio motivation, punishment resistance, and fentanyl choice against an alternative reward were not assessed, so measures of fentanyl seeking described here cannot necessarily be equated with compulsive fentanyl use. Finally, group sizes were larger for C57BL/6 than for CD1 mice and were not explicitly designed for targeting sex differences, so the sparse and inconsistent sex effects reported here should be treated as informative but not conclusive.

Rather than identifying a single mouse strain as the optimal model of fentanyl use disorder, these results indicate that choice of strain is investigation dependent. Strain of mice should be selected according to the specific component of fentanyl vulnerability being studied. Given the continuous distribution of severity scores within strains, the model also provides a framework for testing whether neural adaptations predict or emerge from patterns of fentanyl-use risk. Detailed protocols and standardized analysis code are openly available to support adoption across laboratories.

## Conflict of interest statement

The authors declare no competing financial interests.

## Funding statement

This project was funded by grants from the National Institute of Health (NIDA R00-DA052571 to K.R.C, R01-DA059374 to S.A.G. and J.F.N., NEI T32-EY007031 to T.R.A., NINDS R25-NS114097 to E.S.)

## Ethics approval statement

Experimental procedures were approved by the University of Washington Institutional Animal Care and Use Committee (IACUC).

---

**Figure S1.**
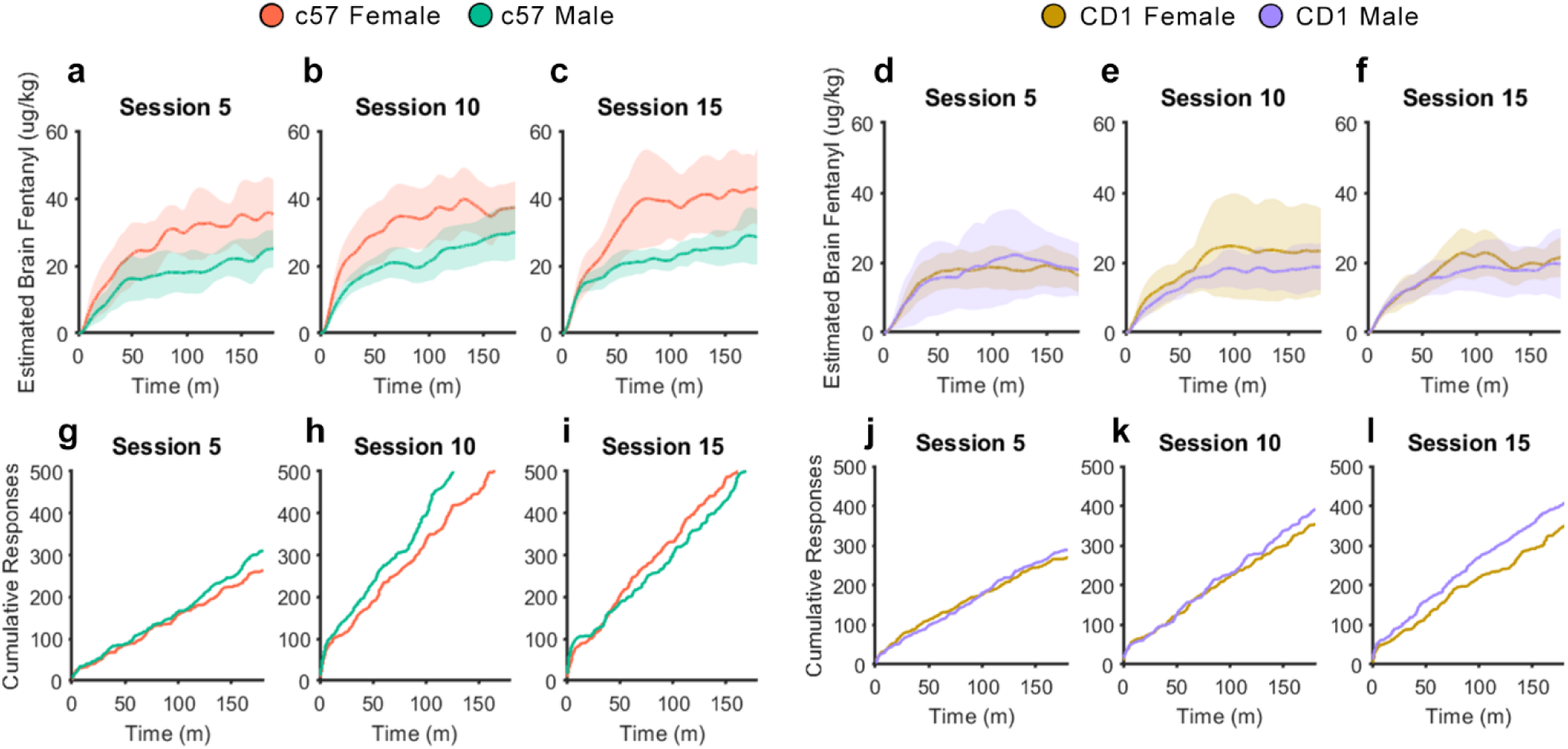
Estimated Brain Fentanyl Levels. Average estimated brain fentanyl concentration within session for c57 mice on **a)** session 5, **b)** session 10, and **c)** session 15. Average estimated brain fentanyl concentration within session for CD1 mice on **d)** session 5, **e)** session 10, and **f)** session 15. Cumulative active lever presses for c57 mice on **g)** session 5, **h)** session 10, and **i)** session 15. Cumulative active lever presses for CD1 mice on **j)** session 5, **k)** session 10, and **l)** session 15.

## Notes

### Competing Interest Statement

The authors have declared no competing interest.

